# The effects of experimental floral resource removal on plant-pollinator interactions

**DOI:** 10.1101/2021.03.21.436328

**Authors:** Justin A. Bain, Rachel G. Dickson, Andrea M. Gruver, Paul J. CaraDonna

## Abstract

Pollination is essential for ecosystem functioning, yet our understanding of the empirical consequences of species loss for plant-pollinator interactions remains limited. It is hypothesized that the loss of abundant and generalized (well-connected) species from a pollination network will have a large effect on the remaining species and their interactions. However, to date, relatively few studies have experimentally removed species from their natural setting to address this hypothesis. We investigated the consequences of losing an abundant, well-linked species from a series of plant-pollinator networks by experimentally removing the flowers of *Helianthella quinquenervis* (Asteraceae) from half of a series of 10 paired plots (15 m diameter) within a subalpine ecosystem. We then asked how the localized loss of this species influenced pollinator visitation patterns, floral visitor composition, and interaction network structure. The experimental removal of *Helianthella* flowers led to an overall decline in plot-level pollinator visitation rates and shifts in pollinator composition. Species-level responses to floral removal differed between the two other abundant, co-flowering plants in our experiment: *Potentilla pulcherrima* received higher visitation rates, whereas *Erigeron speciosus* visitation rates did not change. Experimental floral removal altered the structural properties of the localized plant-pollinator networks such that they were more specialized, less nested, and less robust to further species loss. Such changes to interaction structure were consistently driven more by species turnover than by interaction rewiring. Our findings suggest that the local loss of an abundant, well-linked, generalist plant can bring about diverse responses within pollination networks, including potential competitive and facilitative effects for individual species, changes to network structure that may render them more sensitive to future change, but also numerous changes to interactions that may also suggest flexibility in response to species loss.

## Introduction

Plant-pollinator interactions are essential for ecosystem functioning. It is estimated that nearly 90% of flowering plant species depend on animal visitors for some aspect of their own reproduction (Ollerton et al. 2011), and more than 200,000 animal species rely on floral resources for food (Inouye and Ogilvie 2017). Global environmental change—including climate change, habitat loss, and invasive species—is threatening nearly 40% of vascular plant species with extinction globally (Lughadha et al. 2020), and declines in pollinator populations are becoming increasingly documented (Burkle et al. 2013, Cameron and Sadd 2020). Because of their mutualistic relations, the loss or reduction of plants or pollinators can have important consequences for the remaining species, their interactions, and the structure of plant-pollinator networks (e.g., Memmott et al. 2004; Burkle et al. 2013; Mathiasson and Rehan 2020).

Although rare and more specialized species are hypothesized to be the most susceptible to environmental disturbances (Burkle et al. 2013, Mathiasson and Rehan 2020), the loss of a more abundant generalist species from an interaction network may have disproportionately strong effects on the remaining species and their interactions (Memmott et al. 2004). Generalist species are often abundant and well-connected within pollination networks (Waser et al. 1996, Ollerton et al. 2007, Fort et al. 2016). Consequently, disturbances that reduce or remove generalists from a network may bring about considerable and immediate consequences. While this prediction is well supported by simulation models that sequentially eliminate species (and therefore all of their interactions) from networks to understand which species may bring about the largest consequences (e.g., Memmott et al. 2004), it remains less clear how the loss of a well-connected generalist may play out in nature.

To date, relatively few studies have experimentally removed plant or pollinator species from intact pollination networks to investigate the consequences of their loss, but the available evidence suggests a variety of responses (plant removals: Biella et al. 2020; Biella et al., 2019; Ferrero et al. 2013; Goldstein and Zych 2016; Kaiser-Bunbury et al. 2017; pollinator removals: Brosi and Briggs 2013; Brosi et al. 2017). Complete removal of generalist floral resources from a landscape can bring about dramatic reductions in visitation rates and redistribution of visitors to the remaining plant community (Biella et al. 2019). If generalist plants facilitate pollinator visitation (Ghazoul 2006), then the local loss of a generalist plant may increase competition among the remaining flowering plant species for pollinator visitation, with potential consequences for their reproduction (Mitchell et al. 2009). In contrast, if the remaining floral resources are less attractive to pollinators, the loss of a generalist plant may push pollinators to forage elsewhere. Such shifts in pollinator foraging may functionally act as pollinator species loss from a localized interaction network. In concert, these changes may restructure interaction networks such that they become more specialized (e.g., Biella et al. 2020) or more generalized (e.g., Goldstein and Zych 2016; Brosi et al. 2017)—with possible consequences for their sensitivity to future perturbations.

Here, we investigated the empirical consequences of the loss of an abundant, well-linked flowering plant species from an intact plant-pollinator community in the Colorado Rocky Mountains. We did this by experimentally removing the flowers of *Helianthella quinquenervis* (Asteraceae; hereafter *Helianthella*) from half of a series of 10 paired plots (15 m diameter). Our floral removal experiment aims to mimic a natural ecological disturbance that alters the local plant-pollinator landscape in our ecosystem—episodic spring frost events (Inouye 2000). In particular, in high-elevation and high-latitude locations, a consistent pattern of recent climate change is earlier spring snowmelt, which causes plants to begin flowering earlier in the spring (Iler et al. 2013, CaraDonna et al. 2014). This shift in flowering time means that some species are more susceptible to nighttime frost events when the flower buds are developing, including *Helianthella* (which is particularly sensitive to these frost events in our system). As a result, *Helianthella* flower abundance can be dramatically reduced, and in some years there may be no *Helianthella* flowers in entire meadows due to frost damage (Inouye 2008, Iler et al. 2019). Using this experimental setup, we asked how the removal of *Helianthella* flowers influenced: (1) community-level pollinator visitation rates, species-level visitation rates to other co-flowering plant species, and the composition of the pollinator community; (2) the structural properties of the plant-pollinator interaction networks, including whether changes in structure were attributed to interaction rewiring or species turnover. Altogether, this work provides an experimental test to improve our understanding of the consequences of losing an abundant, well-linked, generalist plant species in response to a natural ecological disturbance.

## Materials and Methods

### Study site & study species

Our floral resource removal experiment was conducted in an intact subalpine meadow primarily composed of perennial herbs and bunchgrasses near the Rocky Mountain Biological Laboratory (RMBL) in Gothic, Colorado, USA (38°57.50N, 106°59.30W, 2900 m above sea level). The growing season in this subalpine ecosystem is brief, lasting approximately 3–5 months (CaraDonna et al. 2014). The plant and pollinator communities are relatively generalized and consist almost exclusively of native plants and pollinators (CaraDonna et al. 2017, CaraDonna and Waser 2020). The European honey bee (*Apis mellifera*) is absent near the RMBL. Because non-native honey bees can compete with and influence the behavior of native foraging pollinators (e.g., Thomson 2016, Cane and Tepedino 2017), our study can explore the effects of an experimentally induced floral resource removal on an intact plant-pollinator community without the confounding effects of the presence of honey bees.

At our study site, three co-flowering perennial wildflower species are the most abundant during mid-summer: (i) *Helianthella quinquenervis* (Asteraceae), (ii) *Potentilla pulcherrima* (Rosaceae; hereafter *Potentilla*), and (iii) *Erigeron speciosus* (Asteraceae; hereafter *Erigeron*) (Fig. 1). Together these three species comprise 91.5% of flowers in our plots during the period of study (Appendix S1: Fig. S1). *Helianthella* is self-incompatible with relatively large, yellow composite flowers (capitulum) that are up to 10 cm in diameter, and plants have flower stalks that are 50–150 cm tall (Weber 1952); it provides moderate to high nectar rewards (mean: nectar sugar concentration = 0.661 mg sugar per μl; nectar volume per flower head = 0.218 μl; sugar per flower head = 0.158 mg); *Helianthella* is particularly sensitive to spring frost events (Inouye 2008). *Potentilla* is self-compatible with small, yellow flowers up to 1 cm in diameter and stalks 30–80 cm tall (Aitken 2004); it provides low to moderate nectar rewards (mean: nectar sugar concentration = 0.379 mg sugar per μl; nectar volume per flower = 0.160 μl; sugar per flower = 0.068 mg). *Erigeron* is largely self-incompatible (P.J. CaraDonna, *unpublished data*) with moderate-sized composite flowers measuring 3–5 cm diameter, composed of yellow disc florets and lavender ray florets, with stems 15–80 cm tall; it provides moderate nectar rewards (mean: nectar sugar concentration = 0.471 mg sugar per μl; nectar volume per flower head = 0.227 μl; sugar per flower head = 0.071 mg). Unlike *Helianthella*, both *Potentilla* and *Erigeron* are not particularly sensitive to spring frost events (CaraDonna and Bain 2016, CaraDonna & Bain *unpublished data*). All three species are visited by a diverse, generalized suite of pollinators (e.g., CaraDonna and Waser 2020), including bumble bees, solitary bees, butterflies, moths, flies, and beetles.

**Figure 1.**
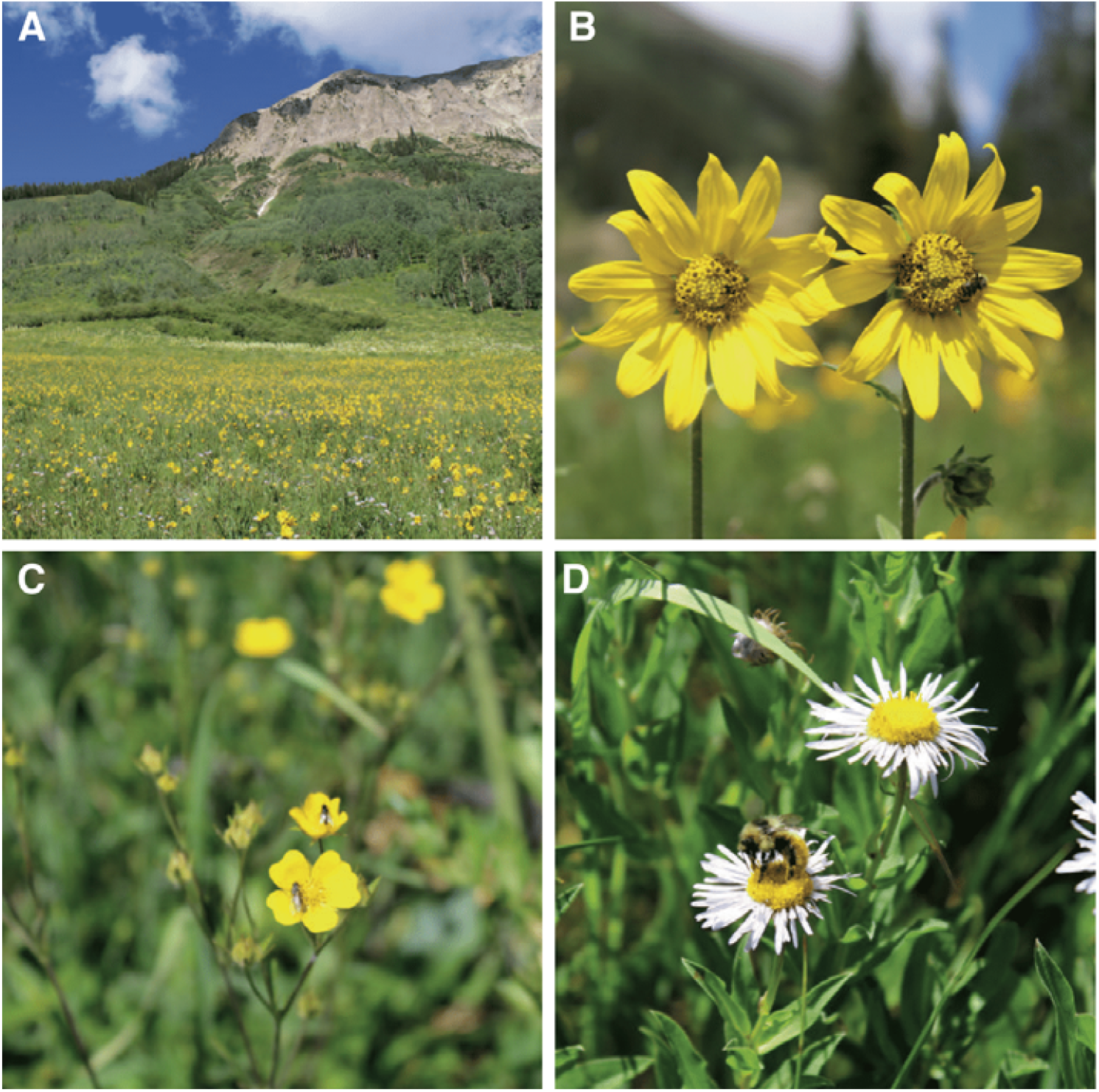
Photographs illustrating (A) the study site at the Rocky Mountain Biological Laboratory in Gothic CO, USA, and (B–C) the three abundant co-flowering plant species. (A) An example of the abundance of *Helianthella quinquenervis* at our study site. (B) *Helianthella quinquenervis*, the focal flowering plant species whose flowers were removed from experimental plots, being visited by a common bumble bee species (*Bombus flavifrons*). (C) *Potentilla pulcherrima* flowers being visited by a common fly species. (D) *Erigeron speciosus* being visited by a common bumble bee species.

### Flower removal experiment

We removed *Helianthella* flowers from a series of large plots at our study site on 5 July 2017 (Fig. 1; Appendix S1: Fig. S2). Our experiment used a paired plot design: 10 *Helianthella* flower removal plots and 10 control plots (*Helianthella* flowers left intact). Each circular plot measured 15 m in diameter (Appendix S1: Fig. S2). Plots were paired based on their spatial proximity within the meadow, whereby the edge of each experimental pair was, on average, 3 m apart. The floral removal treatment was assigned randomly within each plot pair. We cut all *Helianthella* flower heads within floral removal plots while they were in the flower bud stage, approximately one week before the start of *Helianthella* flowering. Any flower heads missed during the initial removal were subsequently removed when they began to bloom. We controlled for any trampling effects that may have occurred during the removal treatment application by replicating a similar trampling effect in all control plots.

Our experimental removal of *Helianthella* flowers aims to mimic a naturally occurring disturbance in our ecosystem—episodic spring frost events—which can effectively reduce or eliminate all *Helianthella* flowers from entire meadows in some years (Inouye 2008, Iler et al. 2019). By observing plant-pollinator interactions and assembling plant-pollinators networks in each of our experimental plots (details below), we explored the effects of the removal of *Helianthella* from a series of replicate, localized interaction networks.

### Plant-pollinator interaction observations

We focused our plant-pollinator interaction observations across the peak flowering period of *Helianthella* over three weeks (10–28 July 2017). Although the entire flowering period of *Helianthella* is somewhat longer than our interaction observations, our sampling period effectively captures pre-peak (week 1), peak (week 2), and post-peak flowering (week 3). Plant-pollinator interactions in control and removal plots were monitored simultaneously by the same two researchers throughout the entire experiment (J.A.B and R.G.D.). We observed each plot twice per observation day: one observation session occurred during the morning (08:30–12:00), and the other occurred during the afternoon (13:00–17:00). During an observation session, we observed each plot for 15 minutes; researchers would slowly move throughout the entire plot, scanning all open flowers a few meters at a time and recording all floral visits by pollinators (*following* CaraDonna et al. 2017). We randomized the order of plot observations before each daily observation session, and researchers swapped treatments after each observation period. We performed all plant-pollinator interaction observations during weather favorable for pollinator activity, when the temperatures were above 15° C, and when it was not raining or excessively windy (< 10 mph). We observed all flowering plant species present during observation sessions to minimize sampling bias attributed to floral abundance. Each plot received a total of six observations per week, resulting in 90 minutes (1.5 hours) of total observation time per plot per week. Across all plots and treatments, this resulted in 30 hours of observation time per week and 90 hours of total observation time across the three-week study period. This sampling level has been shown to effectively characterize the plant-pollinator interactions in this subalpine system (e.g., Burkle and Irwin 2009, CaraDonna et al. 2017).

We recorded an interaction when an insect contacted the reproductive parts of an open flower. For each visitation event, we recorded the plant species visited and the floral visitor’s identity (hereafter referred to as pollinator). We identified pollinators to species, or the lowest taxonomic resolution possible in the field. We did not destructively sample pollinators to avoid influencing plant-pollinator interactions in nearby plots and the following observation period. All plants were identified to species. We quantified floral abundance during each week of the experiment by counting the number of open flowers for all plant species present in a 2 × 2 m square plot, which was located within the center of each larger, circular plot.

### Pollinator visitation and composition

To quantify community-level and species-level pollinator visitation rates, we divided the total number of visits in each plot during each observation session by the total observation time (15 minutes), which provided us with a measure of visits per minute per plot. We accounted for variation in the floral abundance across plots by dividing visitation rates by the total number of flowers recorded in each plot in each week. This calculation provided us with a measure of visits per minute per flower for each plot. We then compared the composition of floral visitors between control and removal plots across all weeks combined using non-metric multidimensional scaling (NMDS) ordination and PERMANOVA. Ordinations were created using square-root-transformed visitation data with Bray-Curtis dissimilarity distances (Oksanen et al. 2019).

### Plant-pollinator interaction network structure

We constructed a series of plant-pollinator interaction networks by aggregating interaction data within each week across each plot and each treatment (one interaction matrix per plot per week). To investigate how the removal of *Helianthella* flowers from our experimental plots influenced the structure of plant-pollinator interaction networks, we compared six measures of network structure between the control and removal treatments: connectance, links per species, network-level specialization, nestedness, network robustness, and plant and pollinator niche overlap. Together, these metrics help reveal whether the structure of the interaction networks became more or less specialized or generalized and whether they are more or less sensitive to future change. *Connectance* describes a basic component of network complexity and is calculated as the proportion of observed links out of all possible links in the network (values range from 0 to 1). *Network-level specialization (H*_*2*_*’)* is a frequency-based metric that characterizes the level of interaction specialization within a bipartite network; it indicates how much niche partitioning there is across all interactions within the entire pollination network (Blüthgen et al. 2006). Values of *H*_*2*_*’* range from 0 to 1, with higher values indicating greater specialization, and therefore less interaction niche overlap among plants and pollinators within the network. *Weighted nestedness* (*wNODF)*, nestedness based on overlap and decreasing fill(Almeida□Neto et al. 2008); describes the extent to which specialist species (those with few links) interact with generalist species (those with many links) (Bascompte et al. 2003), and takes into account interaction frequencies; values range from 0 to 100, where 100 indicates a perfectly nested network.

*Plant and pollinator niche overlap* measures the amount of interaction specialization separately for plants and pollinators within the network; values range from 0 to 1, where 1 indicates perfect niche overlap (greater generalization) among species. *Network robustness* quantifies a network’s sensitivity to the simulated extinction of plants or pollinators (or both; Memmott et al. 2004, Burgos et al. 2007); robustness values range from 0 to 1, where 0 indicates that all species become secondarily extinct after removing the first species (zero robustness, high sensitivity) and 1 indicates that no species become secondarily extinct (maximum robustness, low sensitivity). We calculated network robustness separately for plants and pollinators by simulating the random loss of species from the opposite group.

Finally, we quantified the amount of interaction turnover (i.e., changes in the composition of interactions) between each pair of control and removal networks. Total interaction turnover between a pair of networks can be attributed to two additive components: species turnover (interactions gained or lost because a species is gained or lost) and interaction rewiring (interactions are reassembled because of changes in who is interacting with whom among species that are present) (Poisot et al. 2012). In the context of our experiment, species turnover represents interactions that are lost (or gained) because a species is absent (or present) between a pair of control and removal networks. Interaction turnover values range from 0 to 1, where 0 indicates no differences in the composition of interactions between networks, and 1 indicates complete interaction turnover (no similarity in composition).

### Data Analysis

We analyzed the effect of experimental treatment (floral removal versus control) on all of our response variables using linear mixed-effects models. In our statistical models, we included observation week as an interaction term to investigate whether any treatment effects varied through time. We included plot pair as a random effect in all models to account for the spatial proximity and repeated sampling of our study plots. We first assessed the significance of treatment effects in our statistical models with a type III ANOVA (i.e., to test whether there was a significant treatment × week interaction); if the interaction term was not statistically significant, we then used a type II ANOVA to assess the significance of the main effects. Before analysis, we checked data for heteroscedasticity and normality.

To understand whether the patterns we observed between treatments were the result of additive or synergistic effects of the experimental removal of *Helianthella* flowers, we repeated each analysis described above with the omission of all interactions with *Helianthella* from our control plot data before treatment comparison. We refer to these as “*Helianthella* omission” analyses. If treatment effects vanish when we omit visits to *Helianthella* from the control plot data, this would suggest that the effect is additive (i.e., driven mainly by the loss of *Helianthella* flowers); in contrast, differences between treatments would suggest that the effect is synergistic (i.e., the removal of *Helianthella* flowers has consequences beyond the loss of its flowers). *Helianthella* omission analyses were otherwise identical to those described above.

All data analyses were performed using R version 4.0.4 (R Core Team 2021). Linear mixed-effects models were performed using the *lme4* package (Bates et al. 2014); all network metrics were calculated in the *bipartite* package (v. 2.15) (Dormann et al. 2008); and interaction turnover was calculated using the *betalink* package (v. 2.2.1) (Poisot et al. 2012, Poisot 2019).

## Results

In total, we observed 9,489 pollinator visits to flowers representing 15 plant species and 60 pollinator species. Across all plots, treatments, and observations, visits to *Helianthella, Potentilla*, and *Erigeron* comprised 91% of all observed visitation events. In the unmanipulated control plots, there were a total of 5,479 visitation events across 15 plant species and 49 pollinator species; *Helianthella* received 2,122 visits (39%) from 32 pollinator species, *Potentilla* received 1,530 visits (28%) from 35 pollinator species, and *Erigeron* received 1,452 visits (27%) from 26 pollinator species. In the *Helianthella* floral removal plots, there were a total of 4,010 visitation events across 11 plant species and 48 pollinator species; here, *Potentilla* received 1,824 visits (45%) from 35 pollinator species, and *Erigeron* received 1,703 visits (42%) from 29 pollinator species. *Potentilla* received visits from 43 pollinator species across all plots and treatments, and *Erigeron* received visits from 35 pollinator species.

*How does experimental floral removal influence pollinator visitation rates and composition?* Removing *Helianthella* flowers reduced pollinator visitation rates at the community-level. On average, community-level visitation rates decreased by 33.3% in removal plots compared to control plots (Table 1; Fig. 2a). Our *Helianthella* omission analysis found no difference in visitation rates between control and removal treatments (Table 1; Appendix S1: Fig. S3), indicating that the reduction in visitation rates is primarily an additive effect of the decrease in flowers within removal treatments.

**Table 1.** Results of the effects of experimental floral removal of a dominant floral resource, *Helianthella quinquenervis*, on pollinator visitation and network structure. In all models, ‘plot pair’ is included as a random effect. Statistical significance for main effects was assessed via type II ANOVA when there was not a significant interaction term in the model. “Community-level *Helianthella* omission” indicates analysis where all visits to *Helianthella* are omitted from control plot data before treatment comparison (see Materials and Methods for details).

| Pollinator visitation rates | | $\chi^2$ | *P* |
| --- | --- | --- | --- |
| Community-level | treatment | 15.331 | < 0.001 |
|  | week | 116.36 | < 0.001 |
| | treatment × week | 1.868 | 0.393 |
| Community-level<br>(*Helianthella* omission) | treatment | 0.305 | 0.581 |
|  | week | 67.528 | 0.091 |
| | treatment × week | 0.169 | 0.844 |
| *Potentilla pulcherrima* | treatment | 14.178 | < 0.001 |
|  | week | 7.935 | 0.019 |
| | treatment × week | 0.222 | 0.895 |
| *Erigeron speciosus* | treatment | 2.066 | 0.151 |
|  | week | 82.534 | < 0.001 |
| | treatment × week | 1.656 | 0.198 |
| Network structure |  |  |  |
| Connectance | treatment | 3.295 | 0.07 |
|  | week | 3.461 | 0.177 |
| | treatment × week | 6.722 | 0.035 |
| Specialization ($H_2'$) | treatment | 5.527 | 0.023 |
|  | week | 7.414 | 0.002 |
| | treatment × week | 0.135 | 0.874 |
| Nestedness (wNODF) | treatment | 8.482 | 0.0056 |
|  | week | 6.579 | 0.0031 |
| | treatment × week | 1.51 | 0.232 |
| Niche Overlap (pollinators) | treatment | 16.18 | < 0.001 |
|  | week | 16.81 | < 0.001 |
| | treatment × week | 2.40 | 0.300 |
| Niche Overlap (plants) | treatment | 19.18 | < 0.001 |
|  | week | 5.05 | 0.08 |
| | treatment × week | 15.49 | < 0.001 |
| Robustness (pollinators) | treatment | 11.64 | < 0.001 |
|  | week | 6.165 | 0.046 |
| | treatment × week | 2.554 | 0.279 |
| Robustness (plants) | treatment | 7.617 | 0.0058 |
|  | week | 18.42 | < 0.001 |
| | treatment × week | 2.002 | 0.368 |
| Interaction Turnover | species turnover vs. interaction rewiring | 7.941 | 0.001 |
|  | week | 2.495 | 0.106 |
| | turnover component × week | 0.0181 | 0.982 |

**Figure 2.**
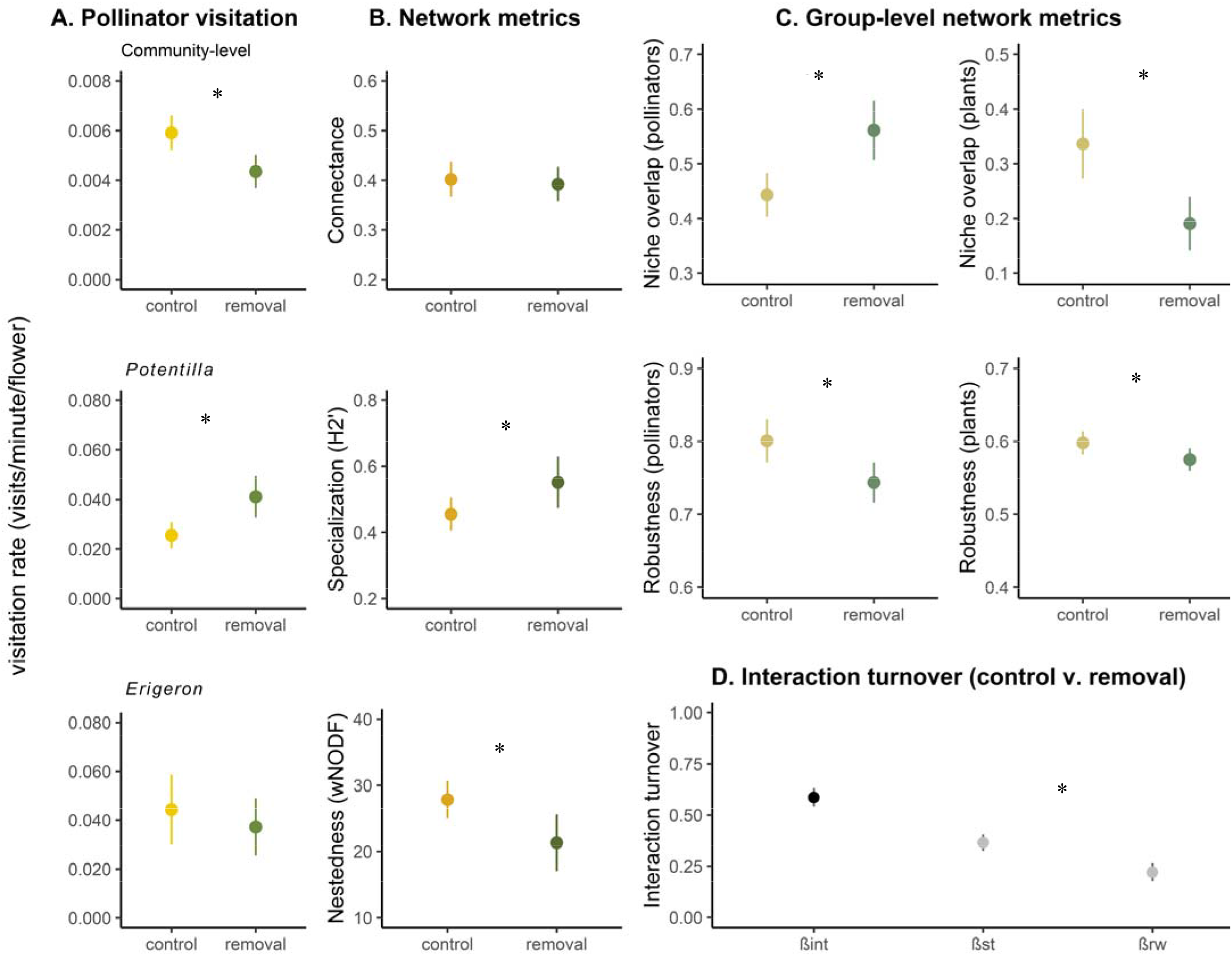
Plots depicting differences between control and the *Helianthella quinquenervis* flower removal treatment for *(a) pollinator visitation* rates at the community-level, to Potentilla pulcherrima, and to Erigeron speciosus; *(b) network metric values* for connectance, specialization (H_2_’), and nestedness (weighted NODF); *(c) group-level metric* values for niche overlap and robustness, and *(d) interaction turnover* where the black dot represents interaction turnover (β_int_) and the gray dots represent species turnover (β_st_) and interaction rewiring (β_rw_) respectively. All dots represent means and error bars represent 95% confidence intervals. * indicates p < 0.05.

Overall, the composition of pollinators differed between control and removal treatments (*F* = 4.33; *P* = 0.002; Appendix S1: Fig. S4). Common bumble bee species contributed most to the species compositional difference between treatments, followed by *Miridae* spp. (leaf bugs), *Rhamphomyia* spp. (dance flies), *Speyeria atlantis* (butterfly) and two solitary bee species, *Halictus virgatellus* and *Megachile melanophaea* (Appendix S1: Table S1). Our *Helianthella* omission analysis found no difference in pollinator composition between the treatments (*F* = 1.52, *P* = 0.12; Appendix S1: Fig. S5), suggesting that the treatment effect was additive and due to the loss of *Helianthella* flowers.

*How does experimental floral removal influence species-level visitation rates to other co-flowering plants?* Removing *Helianthella* flowers resulted in species-specific effects on pollinator visitation rates to the other two abundant co-flowering plant species in the landscape. Visitation rates to *Potentilla* increased by 36.6% on average in floral removal plots compared to control plots (Table 1; Fig. 2a). In contrast, visitation rates to *Erigeron* decreased by 15.9% on average in floral removal plots compared to control plots, but this effect was relatively weak and not statistically significant (Table 1, Fig. 2a).

*How does experimental floral removal influence plant-pollinator network structure?* We found consistent evidence that the removal of *Helianthella* flowers altered the local structure of plant-pollinator interaction networks. All network metrics showed a significant change in response to the removal of *Helianthella*, except for connectance, and such changes were consistent with an overall decrease in interaction generalization (Table 1; Fig. 2b). Network connectance decreased by only 2.5% in the removal networks compared to the control networks, and this pattern was marginally significant (P = 0.07) (Table 1; Fig. 2b). Average network specialization (H_2_’) increased by 17.3% in the removal networks compared to the control networks (Table 1; Fig. 2b). Nestedness (weighted *NODF*) decreased by 23.3% in the removal networks compared to control networks (Table 1; Fig. 2b). There was no significant treatment × week interaction in any metrics, except for connectance, indicating that consistent treatment effects remained even with temporal variation in network structure (Table 1).

Pollinator interaction niche overlap increased by 20.2% in removal networks compared to control networks; in contrast, plant interaction niche overlap decreased by 45.5% in removal networks compared to the control networks (Fig. 2c). We observed significant variation in pollinator niche overlap across weeks but no significant treatment × week interaction; for plant niche overlap, we observed only a significant treatment × week interaction (Table 1).

Network robustness decreased in the floral removal treatment compared to the control networks. Robustness to plant extinctions decreased by 7.5%, whereas robustness to pollinator extinctions by 3.8% (Fig. 2c). We observed significant variation in network robustness across weeks, but we did not observe a significant treatment × week interaction (Table 1).

Our *Helianthella* omission analyses found no difference in network metrics (including group-level metrics) between removal and control networks (except for connectance and plant niche overlap; Appendix S1: Table S2, Fig. S6). This pattern indicates that treatment effects were mostly additive due to the loss of *Helianthella* flowers.

Finally, total interaction turnover between control and removal treatments (β_int_) was high throughout the experiment (mean β_int_ = 0.58) (Fig. 2d). The effect of species turnover (β_st_) on total interaction turnover was greater than the effect of interaction rewiring (β_rw_) (mean β_st_ = 0.36 and mean β_rw_ = 0.22) (Fig. 2d). We did not observe significant variation in interaction turnover across weeks, and we also did not observe a significant treatment × week interaction (Table 1).

## Discussion

Abundant, generalist plant species are highly connected within plant-pollinator networks and interact with many pollinator species (Waser et al. 1996, Vázquez and Aizen 2004, Burkle et al. 2013, Zografou et al. 2020). Disturbances that reduce or eliminate these well-connected species within interaction networks may bring about various consequences for the remaining species and the entire network (e.g., Kaiser□Bunbury et al. 2010, Burkle et al. 2013). Nevertheless, our understanding of the empirical consequences of plant and pollinator species loss remains limited due to the difficulty of removing species under natural conditions in the field (*but see* Brosi and Briggs 2013, Ferrero et al. 2013, Goldstein and Zych 2016, Kaiser-Bunbury et al. 2017, Brosi et al. 2017, Biella et al. 2019, 2020) and in a manner consistent with a natural ecological disturbance. The findings we present here suggest that losing an abundant, generalist floral resource from a localized pollination network may yield numerous changes in interactions among species, including: reduced pollinator visitation rates and altered species composition; potential changes in competition and facilitation among individual plant species; increases in network specialization; and reductions in network robustness.

We experimentally removed *Helianthella* flowers from localized, plant-pollinator networks in a series of large, replicate plots within a subalpine ecosystem. Our experiment aimed to mimic the effects of a natural ecological disturbance—damaging late-spring frost events—that are becoming more frequent in our study system and have clear and dramatic consequences for *Helianthella* (Iler et al. 2019, Inouye 2008). Our results, therefore, represent behavioral changes from pollinators in response to the localized loss of an important floral resource—rather than the result of more extensive extirpation of this species (i.e., from the entire ecosystem)—providing empirical evidence for understanding the consequences of a natural disturbance leading to species loss from interaction networks.

Experimental removal of *Helianthella* flowers resulted in an immediate change in pollinator visitation rates. The observed lower visitation rates in the removal plots appear to be driven mainly by an additive effect of eliminating *Helianthella* flowers, thereby reducing overall plot level floral abundance. Dense and more abundant patches of flowers are generally more attractive to pollinators because they reduce travel time between flowers and increase the foraging efficiency and visitation rates within a patch (Pyke 1984, Sih and Baltus 1987, Ebeling et al. 2008). Evidence points to pollinator visitation frequency as a strong determinant of plant reproductive output (Vázquez et al. 2005). Increased visitation can change the quality of pollination by increasing conspecific pollen deposition, but may also increase rates of heterospecific pollen or self-pollination rates (Karron et al. 1995). Therefore, the reduction in the visitation rates we observed in the *Helianthella* floral removal plots suggests that the absence (or presence) of an abundant and generalized flowering plant has the potential to alter the quantity and quality of pollination services provided by pollinators.

At the species-level, experimental removal of *Helianthella* flowers had contrasting effects on visitation rates for the two other abundant co-flowering plant species in our study. *Potentilla* received increased visitation rates in the absence of *Helianthella* flowers, indicating that it may compete with *Helianthella* for pollinators. A few factors may help to explain these potentially competitive interactions. First, *Helianthella* flowers produce much higher nectar quantities than *Potentilla* (0.158 mg sugar per flower head compared to 0.068 mg sugar per flower), making it an overall higher quality resource to foraging pollinators. Additionally, despite distinct differences in their floral morphology (Fig.1 b and c), both *Helianthella* and *Potentilla* exhibit bright yellow floral displays, which may broadly attract similar assemblages of pollinators, but *Helianthella* flower stalks are much taller than *Potentilla*. Therefore, some pollinators may overlook *Potentilla* when *Helianthella* is present because *Helianthella* flowers are larger, more visible, easily accessible, and provide more floral resources.

In contrast to *Potentilla*, the effect of removing *Helianthella* flowers on visitation rates to *Erigeron* indicated there was either no effect or a potential facilitative relationship. The similarity in floral morphology and visual cues between *Helianthella* and *Erigeron* (Fig.1 b and d) may promote facilitation among plants for pollinators via similar handling strategies and search images (Callaway 1995, Thompson 2001, Johnson et al. 2003, Moeller 2004). *Helianthella* may also act as a magnet species whereby visitors spill over to other nearby, similar, and otherwise less-visited flowers, like *Erigeron* (Laverty 1992, Moeller 2004, Ghazoul 2006). Two traits typically associated with magnet species are large floral displays and the production of large amounts of nectar (Johnson et al. 2003, Molina□Montenegro et al. 2008), both of which *Helianthella* exhibits. Finally, peak flowering time of *Erigeron* occurs after that of *Helianthella* at our study site, and as a result, pollinators initially attracted to *Helianthella* may switch over to later blooming *Erigeron* flowers when *Helianthella* floral abundance starts to decline (Waser and Real 1979). The existence of potentially facilitative interactions between generalists like *Helianthella* and other co-flowering species like *Erigeron* may help to buffer plant and pollinator communities against species extinctions by increasing interaction overlap among pollinators (Verdú and Valiente□Banuet 2008). Consequently, the loss (or reduction) of a generalist like *Helianthella—*and therefore the loss of any facilitative effects—may make plant and pollinator communities more vulnerable to future species losses.

The shift in the floral visitor community in response to the localized experimental removal of *Helianthella* flowers was mainly driven by abundant bumble bee species, which generally avoided the floral removal plots and instead foraged within control plots containing *Helianthella* flowers (Appendix S1: Table S1). Most of the pollinators in our study system have foraging ranges larger than the scale of our experimental floral removal. If *Helianthella* flowers are a preferred resource, then bumble bees and other pollinators can fly to anywhere in the meadow where these flowers are present. This pattern is also consistent with our interaction turnover results, whereby the observed differences in interactions between control and removal networks were primarily driven by changes in species turnover, indicating the movement of pollinators between control and removal treatments. These localized changes in pollinator composition could bring about reproductive consequences for the remaining plant species. For example, bumble bees are the most abundant and effective pollinators in our system, and their shift away from plots without *Helianthella* may lead to a decline in plant reproduction if the remaining floral visitors are less effective pollinators (*sensu* Brosi and Briggs 2013).

The changes we observed in pollinator visitation rates and species composition in response to the removal of *Helianthella* flowers gave rise to the altered structural properties of the plant-pollinator interaction networks. Other experiments that have removed generalist plant species have found conflicting results: Biella et al. (2020) observed increased network specialization, whereas others have observed increased generalization (Goldstein and Zych 2016, Kaiser-Bunbury et al. 2017). Here, we find that experimental floral removal gave rise to more specialized and less nested networks. Such variation in network structure (i.e., the composition of interactions) was due primarily to the effect of species turnover; in other words, the total turnover of interactions between control and removal networks was driven more by interactions lost or gained due to species turnover than it was by interaction rewiring among the remaining species. Overall, the relative differences in network structure between floral removal and control networks remained even in the presence of significant week-to-week variation (which is characteristic of our study system: CaraDonna et al. 2017, CaraDonna and Waser 2020).

More specialized and less nested networks are hypothesized to be less stable for mutualistic networks compared to more generalized and nested networks (Thébault and Fontaine 2010; *but see* Allesina and Tang 2012). Consistent with this hypothesis, when we simulated the random loss of plants or pollinators from control and removal networks, we observed a decrease in robustness in the floral removal networks, indicating that the removal of *Helianthella* flowers at the scale of our experimental plots made the networks more fragile. Robustness decreased almost twice as much for pollinators (in response to simulated plant extinctions) as it did for plants (in response to simulated pollinator extinction), suggesting that the pollinator community may be somewhat more vulnerable to the loss of generalist flowers like *Helianthella* than the plant community to the loss of generalist pollinators. This pattern makes sense given that *Helianthella* is an important food source for many pollinators, but most plant species in this system are visited by multiple pollinators (CaraDonna and Waser 2020). Together, these network properties provide evidence that reducing or removing abundant generalist floral resources, even at a localized, plot-level, may render networks more susceptible to future disturbances.

Although interaction networks became more specialized in response to experimental floral removal, we observed that interaction niche overlap decreased among plants and increased among pollinators when we examined the plants and pollinators as groups separately. In other words, plants became more specialized in response to the local removal of *Helianthella* flowers, whereas pollinators became more generalized. Two factors likely influenced the observed decrease in plant interaction niche overlap: generalist pollinators moving from *Helianthella* removal plots to control plots with *Helianthella* flowers, and a decrease of any potential pollinator facilitation received by plants in the presence of *Helianthella*. Likewise, the *increase* in pollinator interaction niche overlap may also be explained, at least partly, by the decrease of abundant generalist bumble bees in floral removal plots without *Helianthella* flowers. For example, the absence of a highly abundant and generalized pollinator may relax competition among the remaining pollinators, thereby allowing their interaction niches to expand, as was found when dominant bumble bee species were experimentally removed from subalpine meadows (Brosi and Briggs 2013, Brosi et al. 2017). These patterns together suggest variable and flexible responses among plants and pollinator species, which may provide some resilience to the loss (or reduction) of *Helianthella* despite the overall network structure appearing to be at least somewhat more sensitive to further species loss.

Our findings suggest that the removal of an abundant, well-linked, generalist plant from a pollination network can bring about a variety of responses, including potential competitive and facilitative effects for individual species, reductions in network generalization and robustness to species loss, but also numerous changes in interactions among species suggesting some level of flexibility in response to disturbance. By removing *Helianthella* from local plant-pollinator networks, our experiment takes an important step towards understanding how natural ecological disturbances may affect plant-pollinator interactions on a fine temporal and spatial scale. As plant and pollinator declines continue to increase, understanding the consequences of ecological disturbances that reduce or remove abundant and well-connected plant species will be critical.

## Acknowledgments

This work was supported by the Chicago Botanic Garden (P.J.C. and J.A.B.) and an NSF REU fellowship (to R.G.D.). We thank A.M. Iler and K. Mooney for providing access to the study plots; the Rocky Mountain Biological Laboratory for providing access to field sites and facilities; K. Mooney’s lab group, and S. Walwema, for assistance with *Helianthella* removal in the field; R. Perenyi for assistance with flower counts; G. Kirschke for collecting floral nectar data; and the Iler + CaraDonna Lab for constructive feedback on the manuscript.

