## Supporting Information for "The effects of experimental floral resource removal on plant-pollinator interactions"

**Appendix S1**

*The effects of floral resource removal on plant-pollinator interactions: changes in visitation, pollinator composition, and network structure*

***Authors*:** Justin A. Bain[1,2,3], Rachel G. Dickson[3,4], Andrea M. Gruver[1,2], Paul J. CaraDonna[1,2,3]

***Author Institutions:***

[1]Negaunee Institute for Plant Conservation Science and Action, Chicago Botanic Garden, Glencoe, IL, USA

[2]Plant Biology and Conservation Program, Northwestern University, Evanston, IL, USA

[3]Rocky Mountain Biological Laboratory, Crested Butte, CO, USA

[4] Division of Biological Sciences, University of Montana, Missoula, MT, USA

**Table S1**. SIMPER analysis results for the species that contributed most composition differences among the control plots with *Helianthella quinquenervis* flowers and removal plots without *Helianthella* flowers. Species with the lowest values contribute the most to differences among the treatments.

| Species | SIMPER analysis results |
|---|---|
| *Bombus rufocinctus* | 0.120 |
| *Bombus bifarius* | 0.226 |
| *Bombus sylvicola* | 0.315 |
| *Bombus flavifrons* | 0.374 |
| *Halictus virgatellus* | 0.419 |
| *Rhamphomiya spp.* | 0.463 |
| *Bombus appositus* | 0.506 |
| *Bombus californicus* | 0.545 |
| *Speyeria atlantis* | 0.580 |
| *Miridae spp* | 0.611 |
| *Thricops septentrionalis* | 0.641 |
| *Megachile melanophaea* | 0.667 |
| *Tachinidae sp. 1* | 0.691 |
| *Lasioglossum spp* | 0.713 |

**Table S2.** Model coefficients for the analysis of treatment and week on network metrics. In all models, 'plot pair' is included as a random effect. Note that statistical significance for main effects was assessed via type II ANOVA when there was no significant interaction term in the model.

| | | | | ***Helianthella Omission Analyses*** | |
|---|---|---|---|---|---|
| **Network Metric** | | $X^2$ | *P* | $X^2$ | *P* |
| **Connectance** | treatment | 3.295 | 0.07 | 6.414 | 0.01 |
| | week | 3.461 | 0.177 | 3.583 | 0.167 |
| | treatment × week | 6.722 | 0.035 | 3.582 | 0.167 |
| **Specialization ($H_2$')** | treatment | 5.527 | 0.023 | 0.306 | 0.58 |
| | week | 7.414 | 0.002 | 20.72 | < 0.001 |
| | treatment × week | 0.135 | 0.874 | 1.237 | 0.539 |
| **Nestedness (wNODF)** | treatment | 8.482 | 0.0056 | 0.253 | 0.615 |
| | week | 6.579 | 0.0031 | 28.12 | < 0.001 |
| | treatment × week | 1.51 | 0.232 | 4.749 | 0.093 |
| **Niche Overlap (pollinators)** | treatment | 16.18 | < 0.001 | 0.046 | 0.83 |
| | week | 16.81 | < 0.001 | 22.09 | < 0.001 |
| | treatment × week | 2.40 | 0.300 | 0.015 | 0.993 |
| **Niche Overlap (plants)** | treatment | 19.18 | < 0.001 | 5.578 | 0.018 |
| | week | 5.05 | 0.08 | 8.692 | 0.013 |
| | treatment × week | 15.49 | < 0.001 | 5.606 | 0.061 |
| **Robustness (pollinators)** | treatment | 11.64 | < 0.001 | 0.171 | 0.679 |
| | week | 6.165 | 0.046 | 23.50 | < 0.001 |
| | treatment × week | 2.554 | 0.279 | 16.09 | < 0.001 |
| **Robustness (plants)** | treatment | 7.617 | 0.0058 | 0.302 | 0.583 |
| | week | 18.42 | < 0.001 | 16.34 | < 0.001 |
| | treatment × week | 2.002 | 0.368 | 3.273 | 0.155 |

31

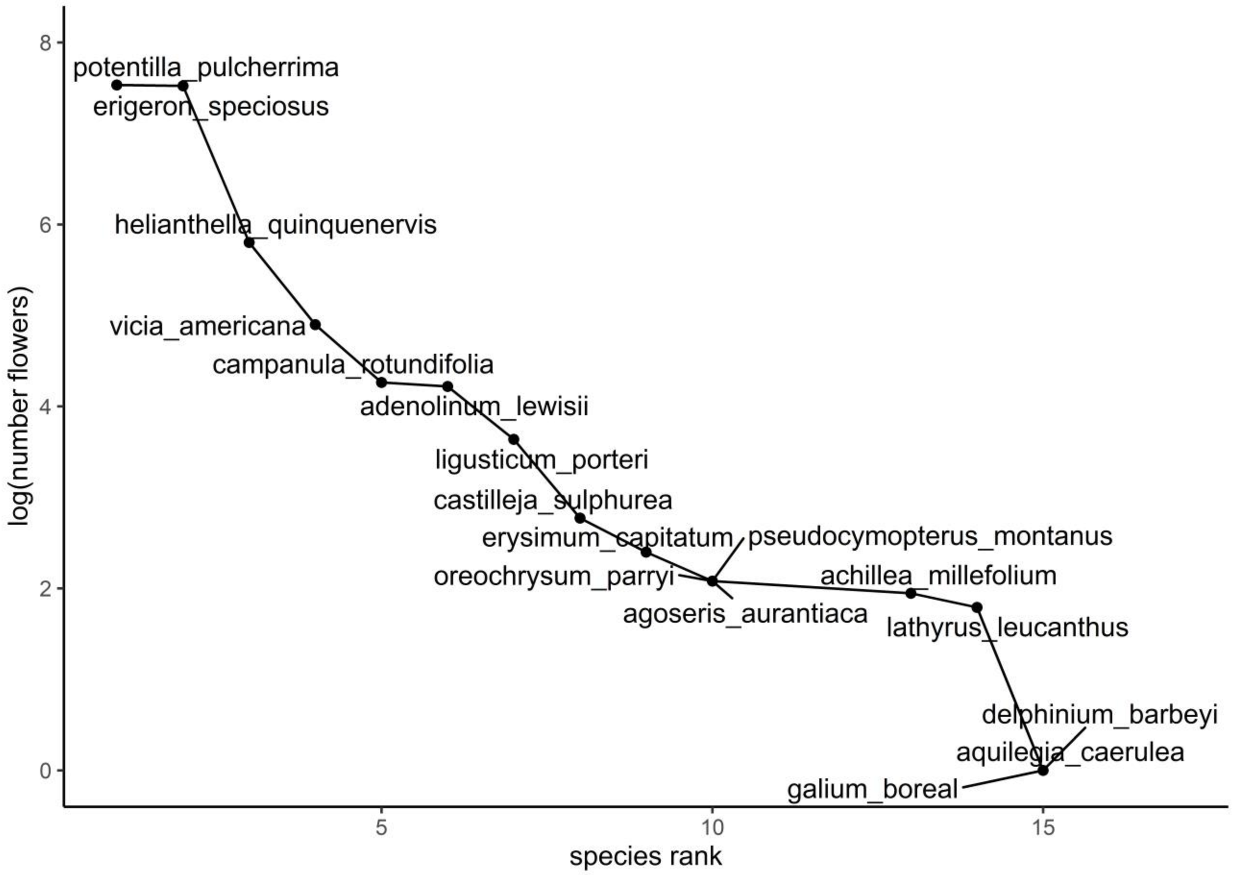


**Figure S1.** Rank abundance curve for all flowering plant species in the plots. The three focal plant species, *Potentilla pulcherrima*, *Erigeron speciosus*, and *Helianthella quinquenervis* were the three most abundant flowers in our plots, in that order.

32

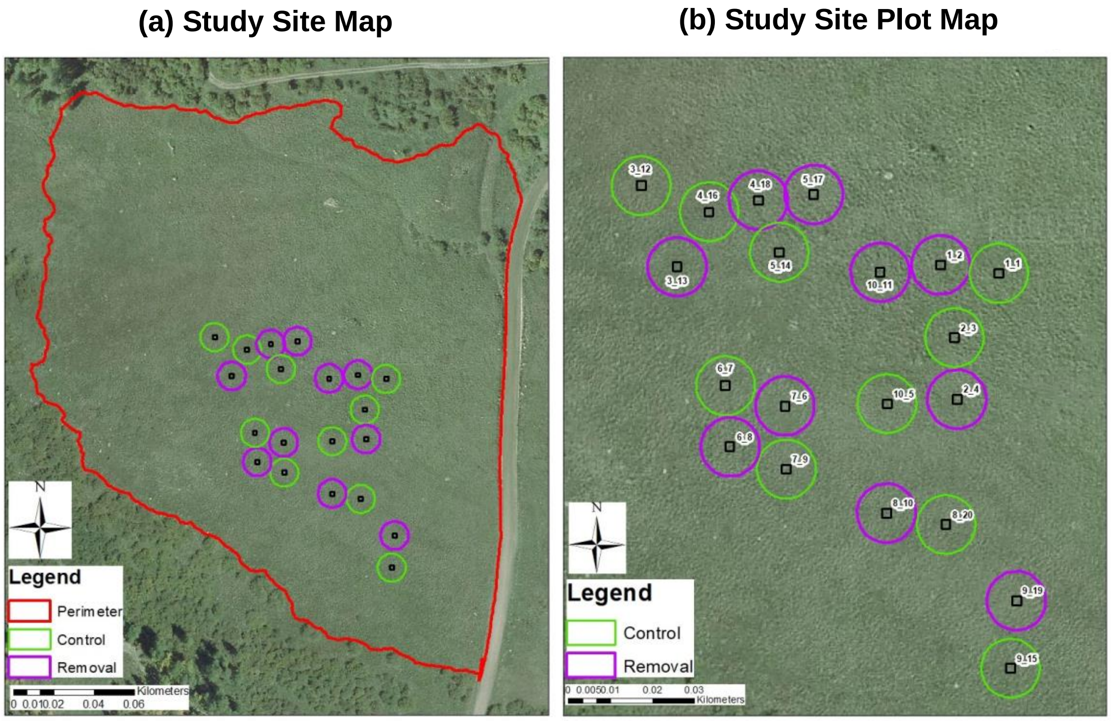


**Figure S2**. (a) Vicinity map of Maxfield Meadow. All green circles are control plots and purple are removal plots. The red line represents the perimeter of Maxfield Meadow. (b) Plot map of Maxfield Meadow. Green circles indicate control plots, and purple indicate removal plots. The first number in the label represents the plot pair number (all removal and control plots are paired) and the second is the individual plot identification number.

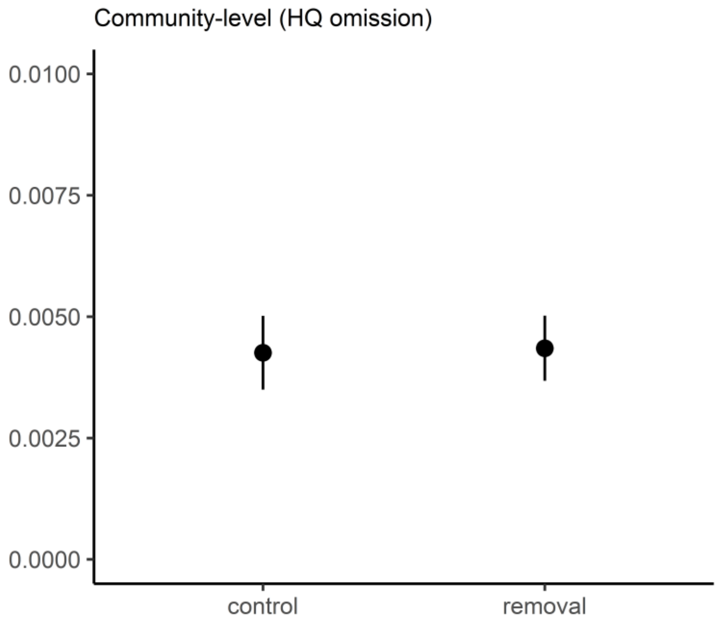


Figure S3. Community-level visitation rates in the control and *Helianthella quinquenervis* flower removal treatment, except interactions with *Helianthella* have been omitted. Dots represent means and error bars represent 95% confidence intervals.

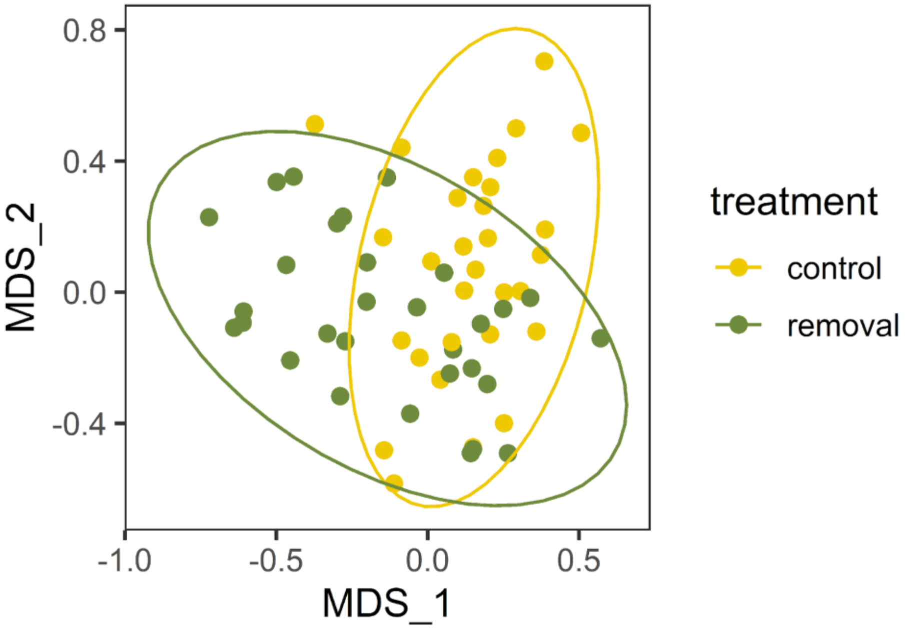


**Figure S4.** Non-metric multidimensional scaling ordination of pollinator communities in overall (i.e., all weeks are pooled and shown here) differences between control treatments with *Helianthella quinquenervis* flowers and removal treatments without *Helianthella* flowers for the entire duration of the experiment (Stress = 0.21; F = 4.33, P = 0.002). The circles represent plots, and the ellipses represent treatments.

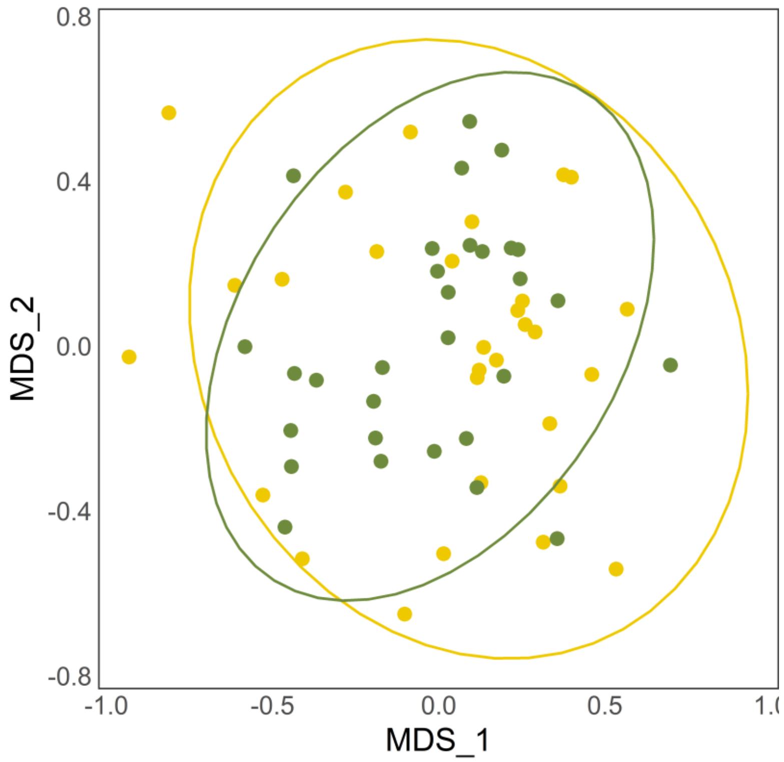


**Figure S5.** Overall (i.e., all weeks are pooled and shown here) non-metric multidimensional scaling ordination of pollinator communities in control and removal plots where the interactions with *Helianthella quinquenervis* have been omitted from the control treatment data (Stress = 0.21; $F = 1.52$, $P = 0.12$). The circles represent plots, and the ellipses represent treatments.

69

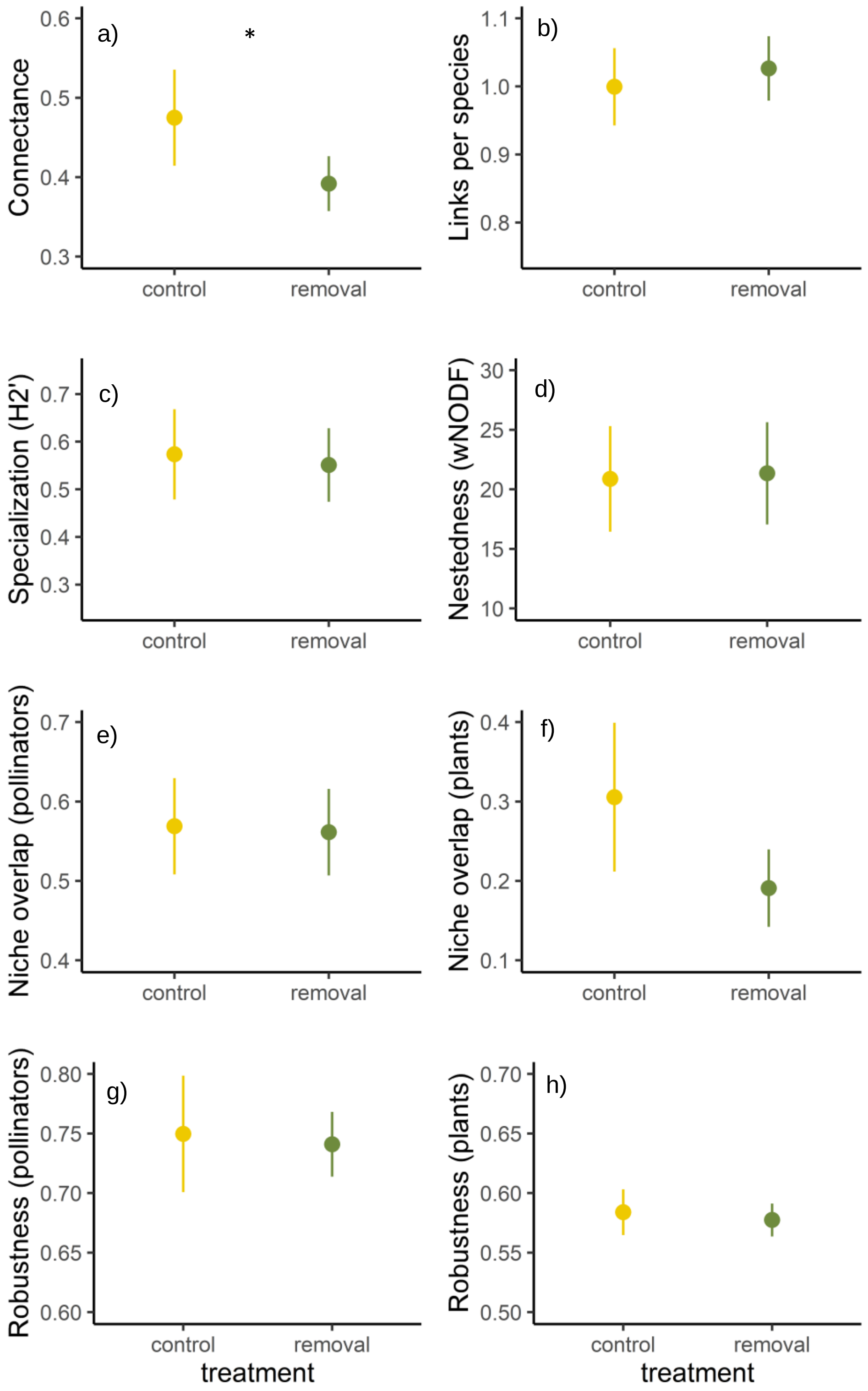
a)
*
Connectance
0.6
0.5
0.4
0.3
control
removal
b)
Links per species
1.1
1.0
0.9
0.8
control
removal
c)
Specialization (H2')
0.7
0.6
0.5
0.4
0.3
control
removal
d)
Nestedness (wNODF)
30
25
20
15
10
control
removal
e)
Niche overlap (pollinators)
0.7
0.6
0.5
0.4
control
removal
f)
Niche overlap (plants)
0.4
0.3
0.2
0.1
control
removal
g)
Robustness (pollinators)
0.80
0.75
0.70
0.65
0.60
control
removal
treatment
h)
Robustness (plants)
0.70
0.65
0.60
0.55
0.50
control
removal
treatment

**Figure S6.** Comparison of network metric values between control and removal treatments but all interactions with *Helianthella quinquenervis* have been removed from the control treatment (*Helianthella* omission analysis): (a) weighted connectance, (b) links per species, (c) Specialization ($H_2$'), (d) weighted nestedness (NODF), (e) Niche overlap (pollinators), (f) niche overlap (plants), (g) Robustness (pollinators), and (h) Robustness (plants). Dots represent means and error bars represent 95% confidence intervals. * indicates $p < 0.05$.
